# High +Gz Exposure Alters Urinary Extracellular Vesicle miRNAs in Humans

**DOI:** 10.64898/2026.11.11.766004

**Authors:** Yusuke Mizohata, Misaki Akiyama, Fumie Takei, Eri Inuzuka, Toru Konishi, Nobuhiro Ohrui

## Abstract

High +Gz exposure elicits complex cardiovascular, autonomic, and endocrine responses, but the associated molecular responses in humans remain poorly understood. We investigated whether microRNAs (miRNAs) encapsulated in urinary extracellular vesicles (uEVs) are altered following acute high +Gz exposure. Twenty-one healthy young men undergoing human centrifuge training provided urine samples before, immediately after, and on the morning after exposure. Candidate uEV-encapsulated miRNAs were identified by RNA sequencing and Gene Ontology analysis, and 16 miRNAs related to ischemia, hypoxia, blood pressure regulation, and mechanical stress were quantified by quantitative PCR. Salivary cortisol, urinary catecholamines, and heart rate were also assessed. Six uEV-encapsulated miRNAs increased significantly after high +Gz exposure. miR-20b-5p, miR-125b-5p, and miR-130a-3p increased immediately after exposure and returned to baseline by the following morning, whereas miR-335-5p, miR-17-5p, and miR-92a-3p remained elevated. Urinary adrenaline, noradrenaline, and dopamine increased immediately after exposure; adrenaline returned to baseline by the following morning, whereas noradrenaline and dopamine remained elevated. Salivary cortisol and heart rate also increased. These findings demonstrate distinct transient and sustained temporal responses of uEV-encapsulated miRNAs to high +Gz in humans, paralleling different catecholamine response patterns. Although their tissue origins and functional roles remain unresolved, these findings provide new molecular-level insights into the human physiological response to high +Gz.

**NEW & NOTEWORTHY:** To our knowledge, this study is the first to demonstrate that acute high +Gz exposure alters miRNAs in human urinary extracellular vesicles. Six miRNAs exhibited two distinct temporal response patterns: three increased transiently, whereas three remained elevated the following morning. These temporal profiles paralleled differential urinary catecholamine responses. The findings reveal a previously unrecognized molecular response to high +Gz and provide new insight into the temporal complexity of human physiological responses to hypergravity stress.

## Introduction

High +Gz exposure (+Gz: acceleration in the head-to-foot direction) is one of the principal aeromedical stressors experienced by fighter pilots and, in severe cases, can culminate in Gravity-induced loss of consciousness (G-LOC) (1). G-LOC represents a serious flight safety risk because it may lead to loss of aircraft control and, consequently, fatal aircraft accidents. High +Gz exposure causes a downward shift of blood toward the lower body and reduces venous return, thereby inducing cerebral hypoperfusion and hypoxia. Consequently, high +Gz exposure elicits multisystem physiological responses involving the cardiovascular, autonomic nervous, and endocrine systems (2). In addition to neuroendocrine activation, +Gz exposure may also induce mechanical stress responses through mechanical forces acting on tissues and cells throughout the body (3, 4).

Acute stress elicits diverse physiological responses that help maintain homeostasis (5, 6). Increases in cortisol and catecholamines, mediated by activation of the hypothalamic–pituitary–adrenal axis (HPA axis) and the sympathetic–adrenal–medullary axis (SAM axis), are key components of the acute stress response; similar neuroendocrine responses have also been reported during acute high +Gz exposure (2). These responses may contribute to the maintenance of cerebral perfusion and systemic homeostasis. However, because high +Gz exposure simultaneously induces hemodynamic alterations, neuroendocrine activation, and mechanical stress, it remains difficult to disentangle the sources and mechanisms of the accompanying molecular changes. Molecular regulators that reflect cellular responses, in addition to transient neuroendocrine changes, may provide insight into these complex physiological responses at the molecular level.

MicroRNAs (miRNAs) are promising candidates for reflecting systemic responses to high +Gz exposure because they post-transcriptionally regulate gene expression and participate in diverse physiological processes, including stress responses, circulatory regulation, and vascular function (7). Moreover, miRNAs can be encapsulated in extracellular vesicles (EVs), which protects them from degradation in body fluids and may allow them to reflect the physiological state of their cells of origin (8–10). Thus, EV-encapsulated miRNAs may serve as useful analytes for noninvasive molecular monitoring.

In this context, analysis of miRNAs encapsulated in urinary EVs may provide a noninvasive approach for assessing systemic molecular responses associated with high +Gz exposure. The urinary EV fraction may include not only EVs derived from the kidney and urinary tract but also molecular components that reflect systemic physiological responses mediated through the circulation (11, 12). Therefore, urinary EV-encapsulated miRNAs may provide a useful window into biological responses to high +Gz exposure.

Beyond providing insight into molecular responses to high +Gz exposure, urinary EV-associated miRNAs may serve as potential biomarkers of physiological adaptation and recovery from gravitational stress. Characterization of these molecular responses may complement conventional physiological and neuroendocrine assessments and advance our understanding of inter-individual variability in responses to high +Gz exposure.

Ultimately, such molecular signatures may facilitate the development of objective strategies for monitoring responses to repeated acceleration stress in aerospace operational settings and may have broader relevance to studies of human adaptation to altered gravitational environments encountered during spaceflight (13).

Accordingly, this study aimed to identify urinary EV-encapsulated miRNAs associated with acute high +Gz exposure and to characterize their temporal response patterns in healthy adults.

## Methods

### Participants

The participants were 21 Japanese male fighter pilot candidates who underwent high +Gz training at the Aviation Safety and Aeromedical Research Group, Japan Air Self-Defense Force, between 23 August 2024, and 22 April 2025 (age, 24.3 ± 1.2 yr) (**Fig. 1A**). Eighteen participants were drug-free, whereas three participants were taking fexofenadine, a medication with no known effect on G tolerance, to manage symptoms caused by allergic rhinitis. (**Fig. 1A**). The study was approved by the Ethical Committee of the Aviation Safety and Aeromedical Research Group (approval number: 5-1-3(1)), and written informed consent was obtained from all participants. All training sessions were conducted under the supervision of a flight surgeon, and all procedures conformed to the relevant guidelines and regulations.

**Fig. 1.**
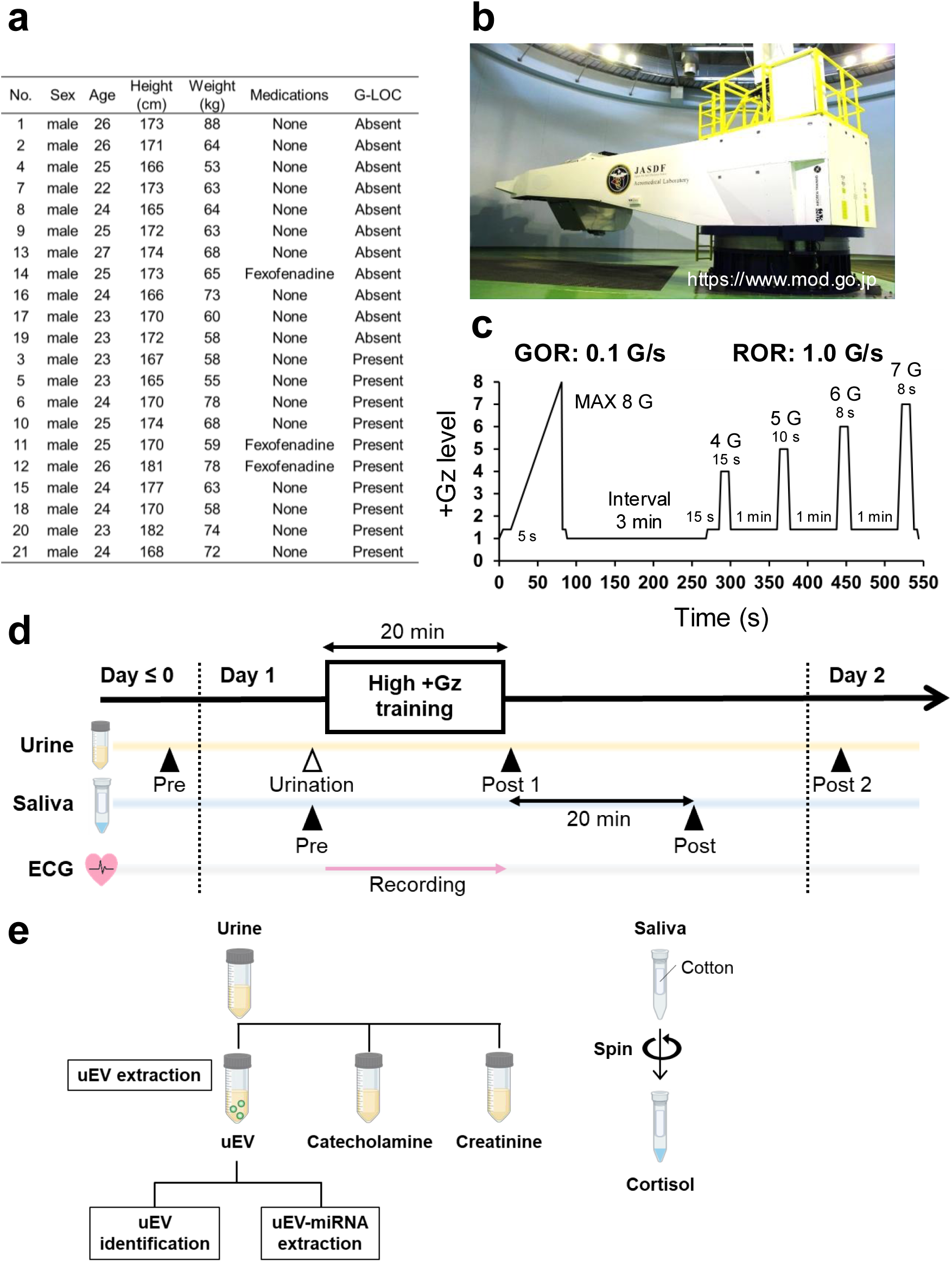
Study design. **a**, Participant characteristics (n=21, 24.3±1.2 years). G-LOC occurred in 10 of 21 participants. **b**, External view of the human centrifuge used for high +Gz training. **c**, High +Gz training profile. Participants were exposed to a gradual onset run (GOR; 0.1 G/s), in which the G-load gradually increased to a maximum of 8 G, and a rapid onset run (ROR; 1.0 G/s) consisting of four exposures to rapidly increasing stepwise +Gz-loads. **d**, Biological sample collection. Urine samples were collected before the day preceding training (Pre), immediately after training (Post1), and on the following day (Post2) (light orange line indicates the urine collection timeline). To assess the effect of high +Gz exposure, participants were instructed to void immediately before training. Saliva samples were collected immediately before training (Pre) and 20 min after training (Post) (light blue line indicates the saliva collection timeline). ECG signals were recorded during high +Gz exposure to monitor heart rate. **e**, Measurements in biological samples. Urine was aliquoted for uEV analysis, catecholamine analysis, and creatinine measurement. Saliva samples were used for cortisol measurement. G-LOC, gravity-induced loss of consciousness. GOR, gradual onset run. ROR, rapid onset run. uEV, urinary extracellular vesicles. miRNA, microRNA.

High +Gz exposure was performed using an ATFS-400J human centrifuge (Environmental Tectonics Corporation, Southampton, PA, United States) located at the Aviation Safety and Aeromedical Research Group (**Fig. 1B**). The high +Gz exposure protocol consisted of a gradual-onset run (GOR; 0.1 G/s), during which +Gz was increased to a maximum of 8 G, followed by a rapid-onset run (ROR; 1 G/s) comprising four stepwise +Gz profiles: 4 G for 15 s, 5 G for 10 s, 6 G for 8 s, and 7 G for 8 s (**Fig. 1C**). After the GOR, participants rested for 3 min before undergoing the ROR. During all training sessions, participants performed the L-1 anti-G straining maneuver (AGSM), which combines repeated straining breathing maneuvers with isometric contraction of the lower-body and abdominal muscles, without anti-G suit support. For GOR, participants initiated the AGSM at the onset of gray-out symptoms, characterized by progressive graying of vision due to reduced retinal perfusion, whereas for ROR, the AGSM was initiated before the onset of +Gz loading. The seatback angle was maintained at 13° throughout all sessions.

Gravity-induced loss of consciousness (G-LOC) was determined by training instructors or a flight surgeon on the basis of signs including loss of visual fixation, eye closure, and inability to maintain upper-body posture during +Gz exposure. When G-LOC occurred, the run was immediately terminated, and +Gz was reduced at an offset rate of 6 G/s. Participants who experienced G-LOC were removed from the centrifuge and provided with sufficient recovery time under the supervision of a flight surgeon. All participants recovered without serious complications.

### Heart rate sampling

Heart rate (HR) was measured during high +Gz exposure (**Fig. 1D**). Electrocardiographic (ECG) signals were recorded using an MP160 system (BIOPAC Systems Inc., Goleta, CA, USA), and instantaneous HR was calculated from the R–R intervals. The instantaneous HR data were then resampled at 0.166-s intervals using cubic spline interpolation. To reduce noise caused by body movements, data points falling outside ±20% of the median value calculated from 25 surrounding beats were considered outliers (14)[**Xu2001JELC**] and excluded from further analysis. The obtained data were compared between the baseline value, defined as the mean value during the 100 s period at 1 G immediately before high +Gz exposure, and the peak value attained during the exposure.

### Urine and saliva collection

Urine samples were collected 4 days before high +Gz exposure between 8:30 and 10:30 (Pre), immediately after exposure (Post1), and between 8:00 and 9:00 on the next morning after exposure (Post2) (**Fig. 1D**). At each sampling point, urine was divided into three aliquots for urinary extracellular vesicle (uEV) extraction, catecholamine measurement, and creatinine measurement (**Fig. 1E**). For uEV extraction, urine was centrifuged to remove cell debris (3,000 rpm, 15 min, at room temperature), and 10 mL of the supernatant was collected. For catecholamine measurement, concentrated hydrochloric acid was added to acidify the urine. For creatinine measurement, aliquoted urine was stored without further treatment. All urine samples were stored at −80°C until analysis.

Saliva samples for cortisol measurement were collected between 8:00 AM and 9:00 AM on the day of high +Gz exposure (Pre) and immediately after exposure, defined as 20 min after termination of the centrifuge run (Post1) (**Fig. 1D**). Saliva was collected using Salivette tubes with cotton swabs (Sarstedt, Nümbrecht, Germany). Participants allowed saliva to be absorbed into the swab according to the manufacturer’s instructions. After centrifugation (3,000 rpm, 15 min, 4°C), the recovered saliva was collected and stored at −80°C until analysis (**Fig. 1E**).

### Identification of Urinary EV-encapsulated microRNAs

uEVs were isolated using a Urine Exosome RNA Isolation Kit (47200; Norgen, Thorold, ON, Canada), which relies on an isoelectric point-based capture method, according to the manufacturer’s instructions (**Fig. 2A**). Briefly, 300 μL of slurry suspension and 250 μL of binding buffer were added to 10 mL of urine supernatant to capture uEVs. After centrifugation (2,500 rpm, 2 min, at room temperature), the resulting pellet was processed for uEV characterization and extraction of encapsulated miRNAs, as described previously (15).

**Fig. 2.**
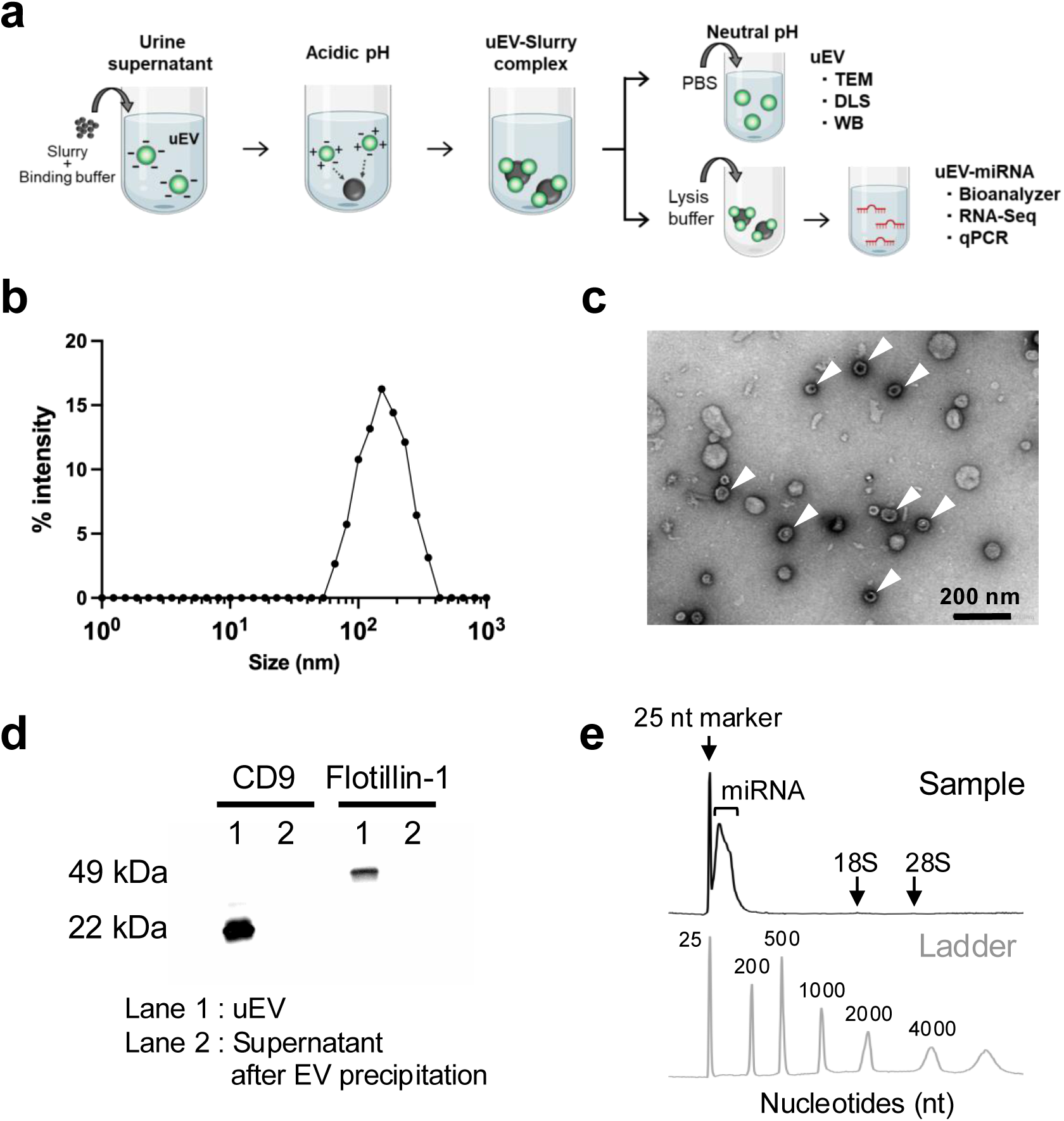
Evaluation of the extracted urinary EVs and miRNAs. **a**, Overview of uEV extraction from urine. **b**,**c**,**d**, Characterization of EVs. In the obtained uEV suspension, DLS showed that particles with sizes characteristic of EVs (50-200 nm) constituted the major population. TEM analysis revealed EV-like morphology (white arrows). CD9 (22 kDa) and Flotillin-1 (49 kDa), EV markers, were detected in the isolated fraction by WB, whereas neither marker was detected in the post-capture urine supernatant. **e**, Electrophoresis analysis. RNA extracted from uEVs showed a distinct enrichment of small RNAs (<200 nt), consistent with miRNAs. In contrast, rRNA peaks were not detected, indicating minimal contamination with cell-derived RNA. uEV, extracellular vesicles. miRNA, microRNA. DLS, dynamic light scattering. TEM, Transmission electron microscopy. WB, Western blotting.

For uEV characterization by transmission electron microscopy (TEM), dynamic light scattering (DLS), and Western blotting, PBS was added to the pellet to release uEVs from the slurry, and the suspension was mixed by vortexing. These methods are recommended by the Minimal Information for Studies of Extracellular Vesicles (MISEV) guidelines and previous studies (16–18). For TEM, the uEV suspension was placed on Formvar-coated copper grids and negatively stained with 3% uranyl acetate. Images were acquired using a JEM-1400plus transmission electron microscope (JEOL, Akishima, Japan) operated at 100 kV. DLS analysis was performed using an ELSZ-1000 system (Otsuka Electronics, Hirakata, Japan) after vortexing to disperse EV aggregates.

For Western blotting, the EV suspension was mixed with 6× Sample Buffer (09499; Nacalai Tesque, Kyoto, Japan) and boiled. After SDS-PAGE with 4 μg of sample per lane and transfer to PVDF membranes, the membranes were blocked with 5% skim milk. The membranes were then incubated with primary antibodies against CD9 (D3H4P Rabbit mAb, 13403S; Cell Signaling Technology, Danvers, MA, United States) and flotillin-1 (D2V7J XP Rabbit mAb, 18634S; Cell Signaling Technology), followed by HRP-conjugated goat anti-rabbit IgG secondary antibody (31460; Thermo Fisher Scientific, Waltham, MA, United States) and ImmunoStar-LD chemiluminescent substrate (290-69904; Wako, Osaka, Japan). Chemiluminescent signals were detected using a ChemiDoc Touch Imaging System (Bio-Rad, Hercules, CA, United States).

miRNAs encapsulated in uEVs were extracted using the same kit according to the manufacturer’s instructions. Briefly, the elution solution supplied with the kit was added to the uEV-bound slurry, and the released miRNAs were bound to a spin column. Finally, miRNAs were eluted from the column using the elution buffer. The quality of the extracted miRNAs was assessed using an RNA 6000 Pico Kit (5067-1513; Agilent Technologies, Santa Clara, CA, United States) on an Agilent 2100 Bioanalyzer. For RNA-seq and qPCR, miRNA concentrations were measured using a Qubit microRNA Assay Kit (Q32880; Invitrogen, Waltham, MA, United States) and a Qubit Flex Fluorometer (Invitrogen), as described previously (19).

### RNA-seq and in silico analysis for Target miRNA selection

To identify miRNAs present in uEVs under normal conditions, uEV-encapsulated miRNAs were extracted from urine samples obtained from two healthy men aged 24 and 25 yr, and comprehensive miRNA-seq analysis was performed by DNA Chip Research Inc. (Tokyo, Japan). miRNAs detected in both individuals were submitted to the miRNA Enrichment Analysis and Annotation Tool (miEAA) (20). Gene Ontology (GO) terms identified in this analysis and considered relevant to high +Gz responses were extracted and scored. The 16 highest-ranked miRNAs based on this scoring process were selected for subsequent qPCR analysis.

### qPCR

Quantitative PCR (qPCR) was performed to identify uEV-encapsulated miRNAs that responded to high +Gz exposure. Reverse transcription was performed using 1 ng of miRNA, a TaqMan^TM^ MicroRNA Reverse Transcription Kit (4366596; Applied Biosystems, Waltham, MA, United States), and 5× RT primers from TaqMan^TM^ MicroRNA Assays (4427975; Applied Biosystems) (**Supplementary Table S1**). The reverse-transcription conditions were 16°C for 30 min, 42°C for 30 min, and 85°C for 5 min. PCR amplification was performed using TaqMan^TM^ Fast Advanced Master Mix for qPCR (4444597; Applied Biosystems) and 20× TaqMan^TM^ MicroRNA Assays (4427975; Applied Biosystems) on a 7500 Fast qPCR System (Applied Biosystems). The PCR conditions were 95°C for 20 s, followed by 40 cycles at 95°C for 3 s and 60°C for 30 s. Relative expression levels were calculated using the 2^−ΔΔCt^ method (21), with the Pre ΔCt value used as the reference. miR-16-5p (22, 23) was used as the endogenous control for ΔCt calculation.

### ELISA for salivary cortisol measurement

Salivary cortisol concentrations were measured using a DetectX Cortisol ELISA Kit (K003-H1; Arbor Assays, Ann Arbor, MI, United States) according to the manufacturer’s instructions. As a preliminary assessment, diurnal variation in salivary cortisol was examined to determine the appropriate timing of saliva collection. Salivary cortisol was measured in two healthy men at seven time points: 5:30, 6:00, 6:30, 8:00, 11:00, 14:00, and 16:30. The results showed a cortisol awakening response and a slight postprandial increase after lunch (**Supplementary Figure S1**). Therefore, pre-exposure saliva samples were collected after 8:00 in participants scheduled for morning high +Gz exposure to avoid the cortisol awakening response and after 13:00 in those scheduled for afternoon exposure to consider the postprandial increase.

### Urinary catecholamine measurement by LC-MS/MS

Urine samples were pretreated before liquid chromatography-tandem mass spectrometry (LC-MS/MS) analysis according to a previously reported (24) method with minor modifications, as follows. After centrifugation (3,600 rpm, 30 min, at room temperature), 300 μL of urine supernatant was mixed with 10 μL deuterated internal standard solution containing 0.05 μg each of adrenaline-D6 ((±)-epinephrine-D6; E-077-1mL; Merck, Darmstadt, Germany), noradrenaline-D6 (norepinephrine hydrochloride-D6; 10842; Cayman Chemical, Ann Arbor, MI, United States), and dopamine-D4 (dopamine hydrochloride-D4; D533782; Fujifilm Wako Chemicals, Osaka, Japan). The sample solution was then heated at 100°C for 20 min to hydrolyze sulfate conjugates, after which 300 μL of 0.05 M ammonium acetate was added for neutralization.

Subsequently, 610 μL of the sample solution was loaded onto an EVOLUTE EXPRESS WCX pretreatment column (602-0001-AXG; Biotage, Uppsala, Sweden) pre-equilibrated with 500 μL of 0.05 M ammonium acetate. After washing with 500 μL of 0.05 M ammonium acetate and 500 μL of 10% methanol, the analytes were eluted with 500 μL of methanol–formic acid (98:2, vol/vol) and evaporated to dryness under a nitrogen stream. The dried samples were reconstituted in 100 μL of 0.1% formic acid so that the final concentration of each deuterated internal standard was 0.5 μg/mL. The reconstituted solution was used for LC-MS/MS injection. Calibration standards were prepared with the same concentrations of deuterated internal standards as those used in the urine samples.

LC-MS/MS analysis was performed using an LCMS-8045 system (Shimadzu, Kyoto, Japan). The injection volume was 5 μL. Chromatographic separation was performed on a Poreshell 120 PFP column (150 × 4.6 mm, 2.7 μm; 693975-408T, Agilent, Santa Clara, CA, United States) at a flow rate of 0.4 mL/min and a column temperature of 40°C. The mobile phases consisted of solvent A, 0.1% aqueous formic acid, and solvent B, acetonitrile. The solvent program was run over a total of 11 min under isocratic conditions at each step: 0–3.5 min, 1% solvent B; 3.5–5.5 min, 98.5% solvent B; and 5.5–11 min, 1% solvent B.

Mass spectrometric detection was performed using the following conditions. The data acquisition window was set at 2.0–3.85 min. The monitored ion transitions (m/z) were 184.20 > 166.05 for adrenaline, 170.20 > 152.10 for noradrenaline, and 154.10 > 137.05 for dopamine. The corresponding transitions for the internal standards were 190.10 > 81.20 for adrenaline-D6, 176.10 > 111.00 for noradrenaline-D6, and 158.10 > 95.10 for dopamine-D4.

### Urinary creatinine measurement by LC-MS/MS

Urine samples were pretreated as follows. 1 μg of Creatinine-D3 (C781502, TRC, Toronto, ON, Canada) was added to 5 μL of urine filtered through a 0.22-μm filter, and the sample was diluted 200-fold with ultrapure water. The diluted solution was used for LC-MS/MS injection.

LC-MS/MS analysis was performed as follows using the system described above (25). The injection volume was 1 μL. Chromatographic separation was performed on a Poroshell 120 EC-C18 column (150 × 2.1 mm, 2.7 μm; 693775-902, Agilent) at a flow rate of 0.3 mL/min. The mobile phases consisted of solvent A, 0.1% aqueous formic acid, and solvent B, acetonitrile. The solvent program was as follows: solvent B was held isocratically at 50% from 0 to 3 min, increased to 98% from 3 to 8 min, and returned to 50% from 8 to 12 min.

Mass spectrometric detection was performed using the following conditions. The data acquisition window was set at 0.5–1.5 min. The monitored ion transitions (m/z) were 114.12 > 44.05 for creatinine and 117.00 > 47.15 for creatinine-D3.

### Statistical analysis

All statistical analyses were performed using GraphPad Prism 10.6.0 (GraphPad Software Inc., San Diego, CA, United States). For comparisons between two groups, unpaired t-tests were used. For multi-group comparisons, one-way or two-way ANO VA was applied followed by Tukey post hoc tests. All tests were two-tailed and set at a significance level of 0.05.

## Results

### Characterization of the isolated urinary EVs and miRNAs

The isolated uEVs were characterized by TEM, DLS, and Western blotting, as recommended by MISEV2023 (16) and previous reports (17, 18). DLS analysis demonstrated that the isolated fraction predominantly contained particles of 100–200 nm, a size range consistent with EVs (**Fig. 2B and Supplementary table S2**), and TEM analysis showed particles with EV-like morphology (**Fig. 2C**). Furthermore, CD9 and flotillin-1, EV markers, were detected in the isolated fraction by Western blotting, whereas neither marker was detected in the post-capture urine supernatant (**Fig. 2D**). These findings indicate that the nano-sized vesicles isolated in this study were consistent with EVs.

Quality assessment of miRNAs extracted from these EVs showed the presence of short RNAs of <200 nucleotides and the absence of cell-derived rRNA. These results confirmed successful recovery of small RNA-enriched miRNA fractions using this method (**Fig. 2E**).

### G-LOC rate during high +Gz training

G-LOC was observed in 10 of the 21 participants (**Fig. 1A**). The maximum +Gz level achieved did not differ between the G-LOC and non-G-LOC groups (**Supplementary Figure S2**).

### Alterations in uEV-encapsulated miRNAs

To identify miRNAs present in uEVs under baseline conditions, RNA-seq analysis was performed. A total of 156 miRNAs were detected in both individuals (**Fig. 3A and Supplementary Table S3**). GO analysis of these 156 miRNAs using miEAA identified 36 GO terms (**Supplementary Table S4**). Because high +Gz exposure is associated with cerebral ischemia/hypoxia, increased blood pressure, and mechanical stress responses, GO terms related to these processes were selected from the 36 identified terms, yielding 14 GO terms.

**Fig. 3.**
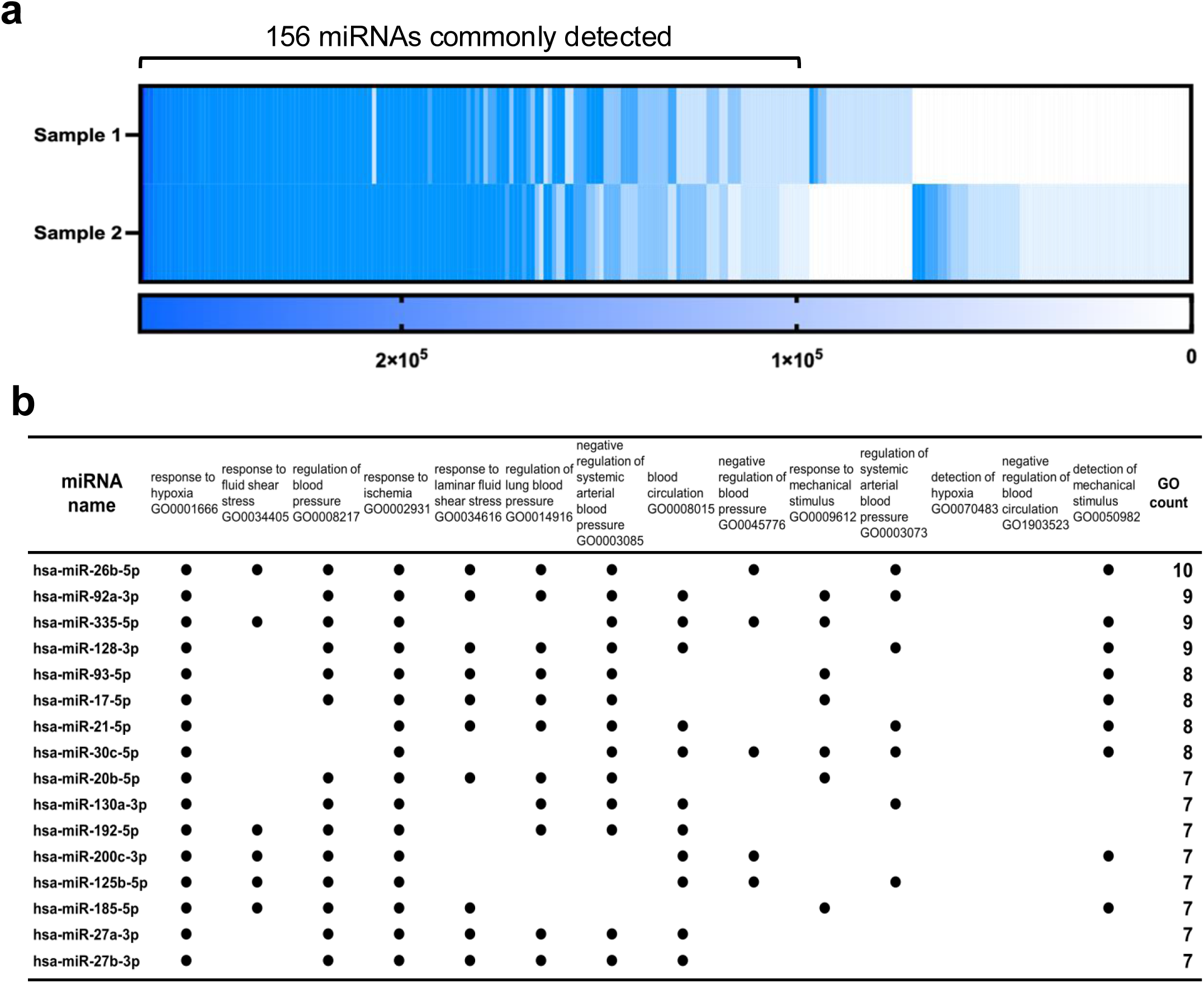
Selection of miRNAs for analysis. **a**, RNA-seq profiling of miRNAs encapsulated in uEVs under the pre-exposure condition. To identify miRNAs present in uEVs under normal conditions, comprehensive RNA-seq profiling of uEV-encapsulated miRNAs was performed using samples from two individuals without +Gz exposure. A total of 156 miRNAs were commonly detected in both samples. **b**, Selection of candidate miRNAs potentially involved in the high-G response. From the 156 miRNAs identified in ***a***, miEAA analysis was performed to extract Gene Ontology (GO) terms related to hypoxia, mechanical stress, and blood pressure. miRNAs were ranked according to the number of associated GO terms, and 16 miRNAs with a total GO count ≥7 were selected for qPCR analysis. miRNA, microRNA.

For each of the 156 miRNAs, the number of associated GO terms among the 14 selected terms was counted. The miRNAs were then ranked in descending order according to this count. The 16 highest-ranked miRNAs were miR-26b-5p, miR-92a-3p, miR-335-5p, miR-128-5p, miR-93-5p, miR-17-5p, miR-21-5p, miR-30c-5p, miR-20b-5p, miR-130a-3p, miR-192-5p, miR-200c-3p, miR-125b-5p, miR-185-5p, miR-27a-3p, and miR-27b-3p (**Fig. 3B** and **Supplementary Table S4**).

These 16 miRNAs were therefore analyzed for changes after high +Gz exposure. The total amount of uEV-encapsulated miRNAs was not significantly altered by high +Gz exposure. (**Supplementary Figure S3**). miR-20b-5p, miR-125b-5p, and miR-130a-3p increased significantly at Post1 and returned to baseline levels at Post2 (**Fig. 4A and Supplementary table S5**). In contrast, miR-335-5p, miR-17-5p, and miR-92a-3p increased significantly at Post1 and remained elevated at Post2 (**Fig. 4A and Supplementary table S5**). The levels of the remaining 10 miRNAs were not significantly altered by high +Gz exposure (**Fig. 4A and Supplementary table S5**). Fexofenadine use did not affect the changes in these miRNAs (**Supplementary Figure S4**).

**Fig. 4.**
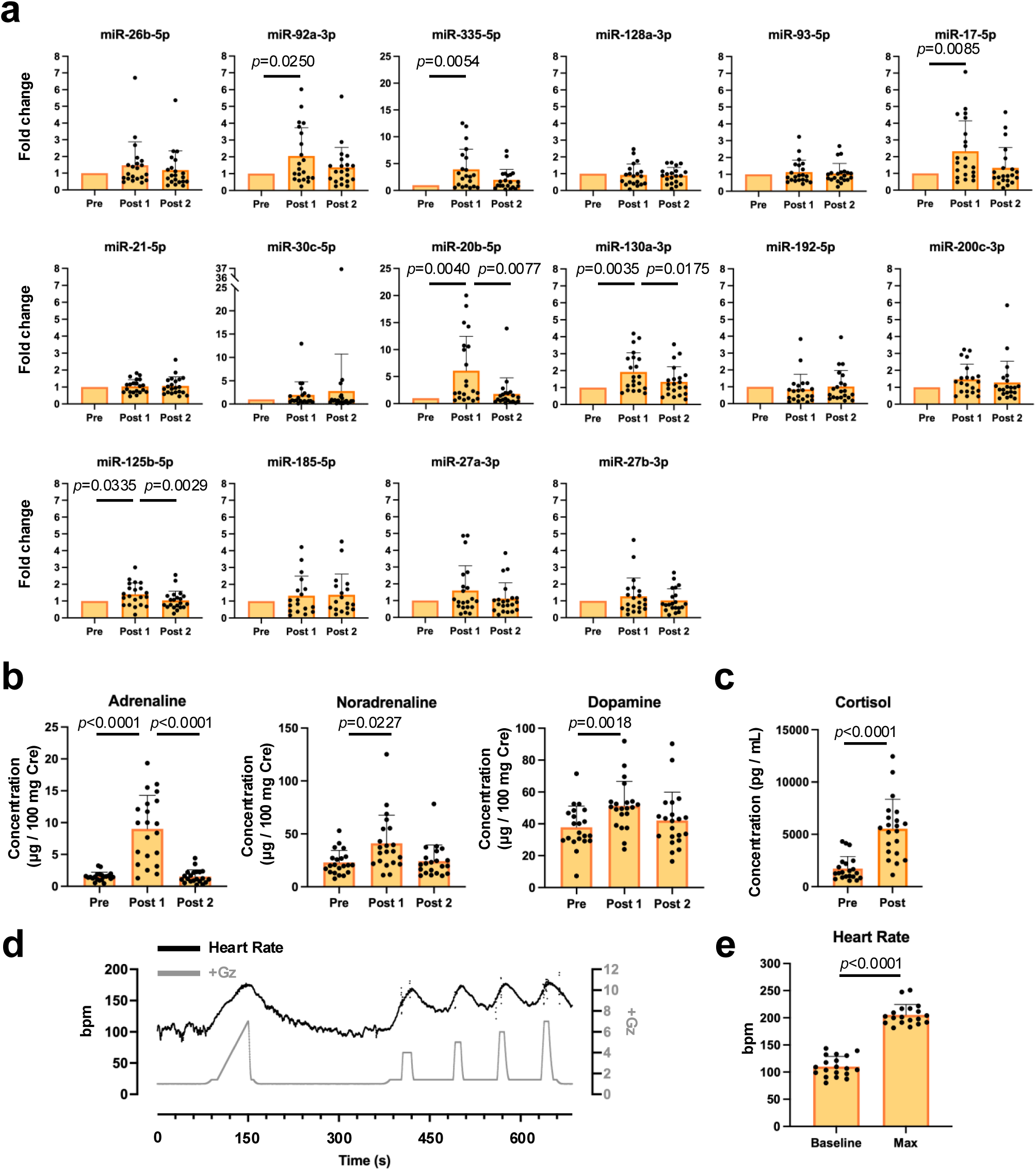
High +Gz stress increases uEV-encapsulated miRNAs and anti-stress reactions. **a**, Alterations in miRNA expression induced by high +Gz stress. Three miRNAs (miR-20b-5p, miR-130a-3p, and miR-125b-5p) significantly increased immediately after +Gz exposure and returned to baseline levels the following day. In contrast, three other miRNAs (miR-92a-3p, miR-335-5p, and miR-17-5p) also increased immediately after +Gz exposure but did not return to baseline levels thereafter. These quantifications were calculated using miR-16-5p as the internal control. **b**, High +Gz stress increased catecholamine level in urine. **c**, High +Gz stress increased cortisol level in saliva. **d**, Representative heart rate recording during +Gz exposure. **e**, High +Gz stress increased heart rate. Data are means±SD. Statistical analysis was performed using one-way ANOVA with the Tukey post hoc test and Student’s two-tailed *t*-test.

### Traditional stress markers

To confirm whether the participants exhibited physiological stress responses, salivary cortisol and urinary catecholamines were measured. Salivary cortisol levels increased significantly after high +Gz exposure (**Fig. 4C**). Similarly, urinary adrenaline, noradrenaline, and dopamine levels increased significantly immediately after high +Gz exposure. Although adrenaline returned to baseline levels by the following day, noradrenaline and dopamine remained elevated (**Fig. 4B**).

### Heart rate

Electrocardiography (ECG) was performed to monitor changes in heart rate during hypergravity exposure. Heart rate increased in accordance with the high +Gz exposure profile (**Fig. 4D**). Comparison of the maximum heart rate during +Gz exposure with the baseline value revealed a significant increase (**Fig. 4E**).

### Effects of G-LOC

To examine whether changes in the six miRNAs increased by high +Gz exposure, salivary cortisol, urinary catecholamines, and heart rate differed according to G-LOC status, participants were stratified into G-LOC and non-G-LOC groups. No significant between-group differences were observed for any of the parameters examined (**Supplementary Figure S5**).

## Discussion

In the present study, we identified six miRNAs (miR-20b-5p, miR-125b-5p, miR-130a-3p, miR-335-5p, miR-17-5p, and miR-92a-3p) contained in uEVs that were increased following high +Gz exposure (**Fig. 3D**). To our knowledge, changes in these miRNAs within uEVs following actual human hypergravity exposure have not previously been demonstrated, representing a novel finding of this study. Because these miRNAs were selected based on GO terms related to blood pressure regulation, hypoxic responses, and mechanical stress (**Fig. 3B**), their alteration in response to hypergravity was therefore biologically plausible. Although uEVs are considered to originate predominantly from cells of the kidney and urinary tract (26, 27), they may also reflect molecular information associated with systemic physiological changes (28, 29). Because the tissues and cell types from which the miRNAs originated were not identified in the present study, our findings do not directly demonstrate responses in specific tissues. Nevertheless, they suggest that at least part of the systemic physiological response induced by high +Gz exposure may be captured noninvasively as changes in uEV-encapsulated miRNAs.

Notably, the six miRNAs exhibited distinct temporal response patterns. miR-20b-5p, miR-125b-5p, and miR-130a-3p increased immediately after exposure and returned to pre-exposure levels by the following morning, whereas miR-335-5p, miR-17-5p, and miR-92a-3p increased immediately after exposure and remained elevated the following morning (**Fig. 4A**). Urinary catecholamines also exhibited distinct temporal patterns: adrenaline increased after exposure and returned to baseline by the following morning, whereas noradrenaline and dopamine remained elevated the following morning (**Fig. 4B, C**). The increases in urinary adrenaline and noradrenaline observed in the present study are also consistent with previously reported changes in circulating catecholamines during high +Gz exposure (30, 31). Although urinary dopamine is less commonly used as an index of sympathoadrenal activation because of its substantial renal contribution, its sustained elevation further supports the presence of prolonged catecholamine-related responses following high +Gz exposure. These findings are consistent with activation of the sympathoadrenal system during acute high +Gz exposure and suggest that hypergravity-induced neuroendocrine responses comprise both transient and sustained components. Although adrenaline exhibited a transient response, the persistent elevations of noradrenaline and dopamine suggest that some components of the catecholaminergic response remain activated beyond the immediate recovery period. Taken together, these findings suggest that hypergravity-induced physiological responses comprise components with distinct temporal characteristics. Interestingly, the temporal patterns of the identified miRNAs paralleled those of the catecholamine responses, with miR-20b-5p, miR-125b-5p, and miR-130a-3p showing transient increases similar to adrenaline, whereas miR-335-5p, miR-17-5p, and miR-92a-3p exhibited more sustained responses resembling those of noradrenaline and dopamine. Furthermore, the increases in salivary cortisol and heart rate observed after high +Gz exposure support the notion that the present exposure elicited a systemic stress response involving both the hypothalamic-pituitary-adrenal (HPA) axis and the sympathoadrenal system.

The three miRNAs showing transient increases (miR-20b-5p, miR-130a-3p, and miR-125b-5p) exhibited temporal changes similar to those of urinary adrenaline, suggesting that their changes may be associated with the acute sympathoadrenal medullary response induced by high +Gz exposure. In the present study, the relative increase in adrenaline immediately after high +Gz exposure was greater than that in noradrenaline and was accompanied by an increase in heart rate, suggesting a prominent acute adrenal medullary response (30, 31). These miRNAs have also been reported to change in response to various forms of acute stress. Increased miR-20b-5p has been reported following psychological stress (32), cerebral ischemia (33), and myocardial ischemia (34). Increased miR-125b-5p has been reported following aerobic exercise (35), cerebral ischemia (36, 37), and β-blocker use (38). miR-130a-3p has been reported to increase following myocardial ischemia (39) and to be involved in responses to mechanical stress (40). Thus, miR-20b-5p and miR-125b-5p may reflect a broader acute stress response rather than a response specific to high +Gz. In contrast, because reports of increased miR-130a-3p following acute psychological or exercise stress are limited, its alteration may be more closely associated with hypoxic or mechanical components of high +Gz exposure than with a nonspecific acute stress response.

In contrast, miR-335-5p, miR-17-5p, and miR-92a-3p, which increased immediately after exposure and had not returned to pre-exposure levels by the following morning, exhibited temporal changes similar to those of urinary noradrenaline. Considering that noradrenaline is released predominantly from postganglionic sympathetic nerve terminals, these miRNAs may be associated with the sustained sympathetic response following high +Gz exposure. Indeed, prolonged noradrenaline responses following psychological stress (41) and exercise (42) have been reported, which is not inconsistent with the present findings. With regard to these miRNAs, increased miR-335-5p has been reported following trauma, which represents a complex stressor (43), whereas increases in miR-17-5p (44) and miR-92a-3p (45) have been reported following acute exercise. In addition, miR-335 has been reported to increase heart rate and blood pressure (46), miR-17-5p to increase following hypoxia (47) and myocardial infarction (48), and miR-92a-3p to inhibit vasodilation (49). Thus, similar to the transiently increased miR-20b-5p and miR-125b-5p, these miRNAs may represent components of a broader stress response that includes responses to the circulatory stress characteristic of high +Gz exposure. Nevertheless, the temporal profile of miR-92a-3p observed in the present study differed from that previously reported (45). The delayed recovery of miR-92a-3p observed here contrasts with the rapid recovery of plasma EV-encapsulated miR-92a-3p after its increase following acute exercise (45), raising the possibility that the delayed recovery of this miRNA may represent a characteristic response to hypergravity.

Interestingly, despite the marked physiological consequences of G-LOC, we did not detect differences between participants who experienced G-LOC and those who did not. This finding suggests that the miRNAs identified here may primarily reflect general physiological responses to high +Gz exposure rather than molecular mechanisms specifically associated with susceptibility to G-LOC. Because the present study employed a candidate-miRNA approach based on predefined biological pathways, it remains possible that other miRNAs more closely related to cerebral hypoperfusion, loss of consciousness, or individual G tolerance were not captured. Future comprehensive profiling studies may therefore be required to identify molecular signatures associated with individual susceptibility to G-LOC.

Although the present findings were obtained following a single high +Gz exposure, the observed temporal patterns suggest that uEV-encapsulated miRNAs may reflect not only acute molecular responses to gravitational stress but also processes involved in physiological recovery.

Taken together, these findings suggest that uEV-encapsulated miRNAs may provide a noninvasive means of capturing molecular responses to gravitational stress. While their utility for monitoring adaptation remains to be established, the identification of miRNAs with distinct temporal response profiles supports the feasibility of using uEV-encapsulated molecular signatures to evaluate recovery following repeated high +Gz exposure. Further studies involving repeated or longitudinal exposure paradigms will be required to determine whether these signatures are associated with longer-term physiological adaptation to gravitational stress.

Finally, several limitations of this study should be acknowledged. First, the tissues or cell types from which the individual uEV-encapsulated miRNAs originated were not identified. Because circulating EV-encapsulated miRNAs were also not measured, it remains difficult to determine whether the observed changes primarily reflect responses within the kidney and urinary tract or broader systemic changes. Future studies using tissue- or cell-specific EV markers together with parallel analysis of circulating EVs will be required to address this issue. Second, we did not directly compare high +Gz exposure with other stressors, such as psychological stress, exercise, or hypoxia. Therefore, it remains unclear whether the observed changes in uEV-encapsulated miRNAs are specific to high +Gz. Future studies comparing the same miRNAs across different stress conditions will be needed to distinguish responses specific to high +Gz from nonspecific stress responses. Third, the target tissues and physiological effects of the uEV-encapsulated miRNAs remain unknown. Although the GO analysis and previous studies provide biological plausibility regarding the potential functions of these miRNAs, their physiological significance was not directly examined in the present study. Further in vitro studies using specific cell types will be required to establish their functional roles. Fourth, the study included only a relatively small sample of young men (n = 21), limiting the generalizability of the findings. Future studies should determine whether similar responses are observed in populations with a broader range of sex, race, and age.

In conclusion, acute high +Gz exposure was associated with increases in six uEV-encapsulated miRNAs that exhibited distinct temporal response patterns. These molecular responses occurred alongside changes in cortisol, catecholamines, and heart rate, suggesting that uEV-encapsulated miRNAs reflect components of the systemic physiological response to hypergravity. Although the biological origins and functional significance of these miRNAs remain to be clarified, the present findings demonstrate the feasibility of using uEV-encapsulated miRNAs as noninvasive markers of responses to gravitational stress and provide a foundation for future studies investigating adaptation to aerospace environments.

## Acknowledgements

We thank the crew of the Centrifuge Training Section, Aviation Physiological Training Division, Aviation Safety and Aeromedical Research Group, Japan Air Self-Defense Force, for their assistance with sample collection during high +Gz training.

## Author contributions

Y. Mizohata and N. Ohrui conceived and designed the study. M. Akiyama performed sample collection, qPCR, ELISA, and LC-MS analyses. YM performed the characterization of uEVs. F. Takei performed the RNA-seq analysis. E. Inuzuka assisted with sample collection. NO, EI and T. Konishi supervised the study. YM wrote the first draft of the manuscript. All authors reviewed and edited the manuscript.

## Data availability

All data in this study were included in this article or online contents. RNA-seq data sets were deposited with GEO (GSE 353823).

## Supplementary Information

Supplementary Figures’ PDF file includes: Figure S1 to S5

Supplementary Tables’ Excel file includes: Table S1 to S5

**Fig. S1.**
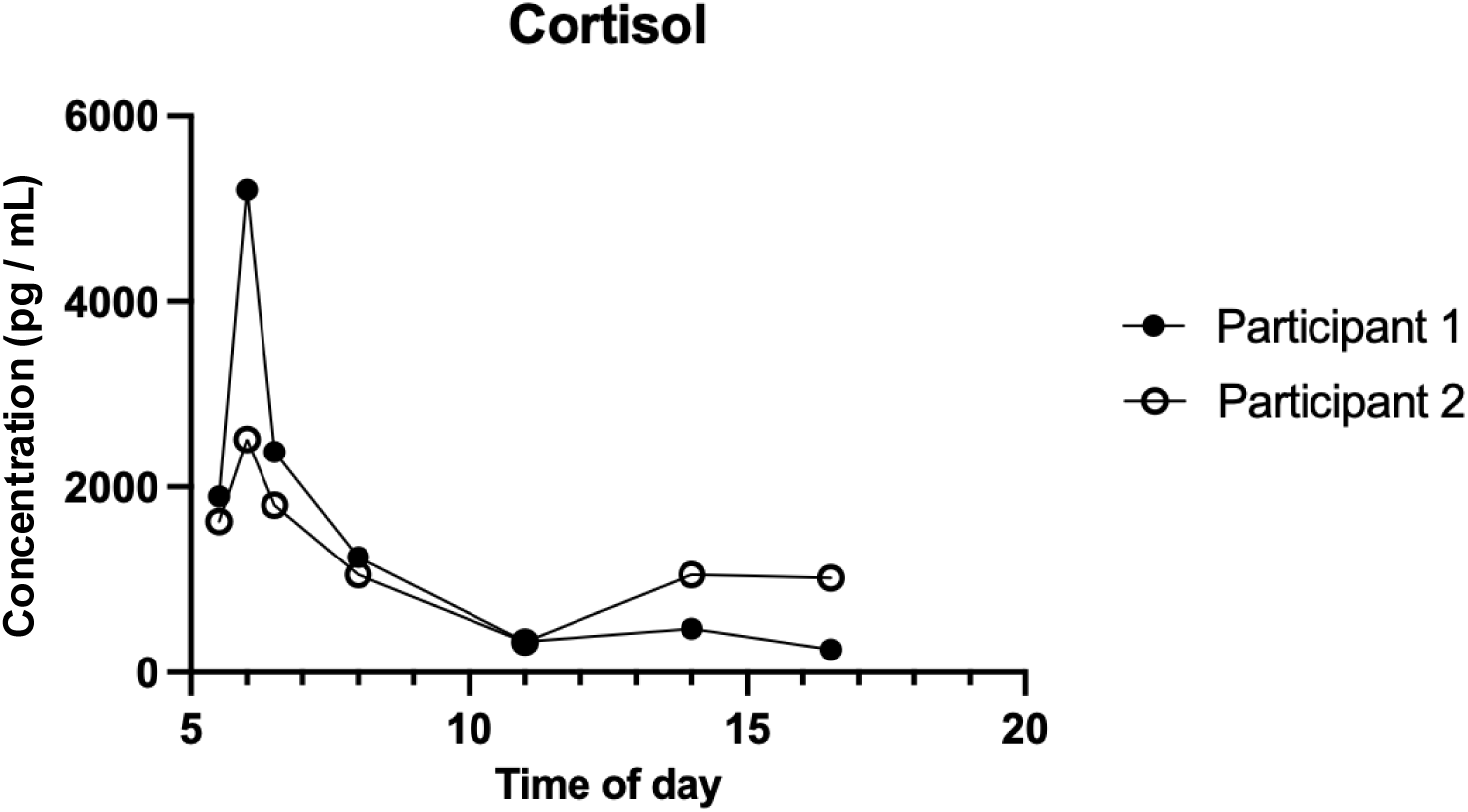
Diurnal variation in salivary cortisol in healthy males. Salivary cortisol exhibited a pronounced increase after awakening (06:00) and a modest increase after lunch (14:00) (n=2).

**Fig. S2.**
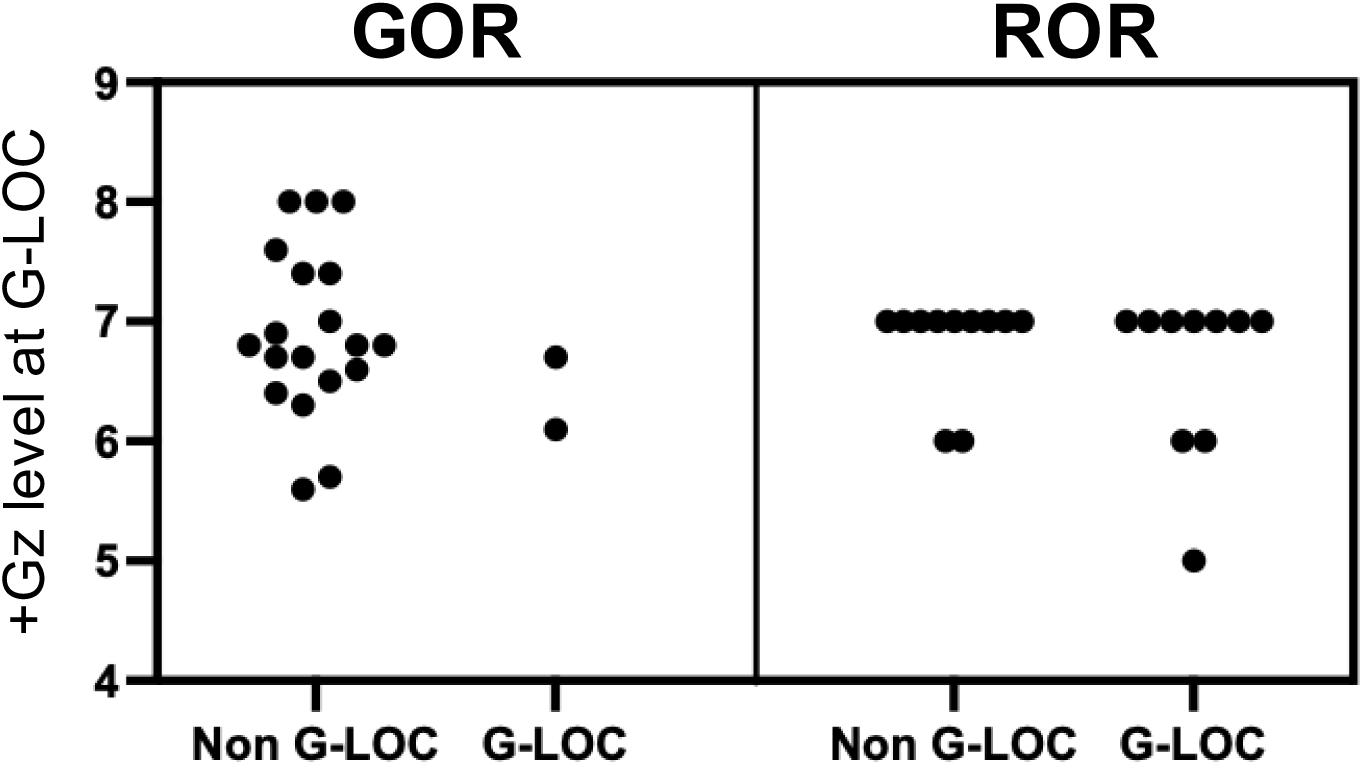
The maximun +Gz levels did not differ between the G-LOC and non-G-LOC groups (n=21). G-LOC, gravity-induced loss of consciousness. GOR, gradual onset run. ROR, rapid onset run.

**Fig. S3.**
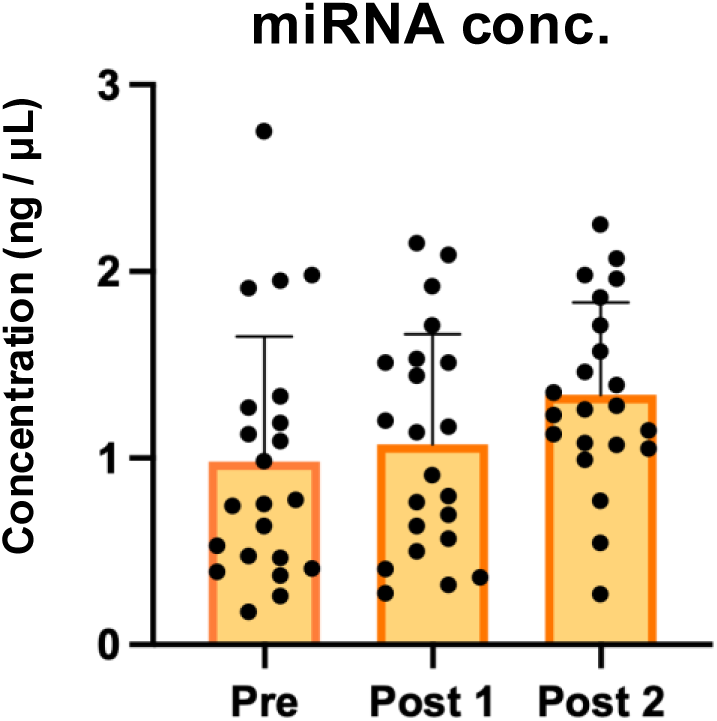
High +Gz stress did not alters amount of uEV-encapsulated miRNAs. Data are means±SD (n=21). uEV, urinary extracellular vesicles. miRNA, microRNA.

**Fig. S4.**
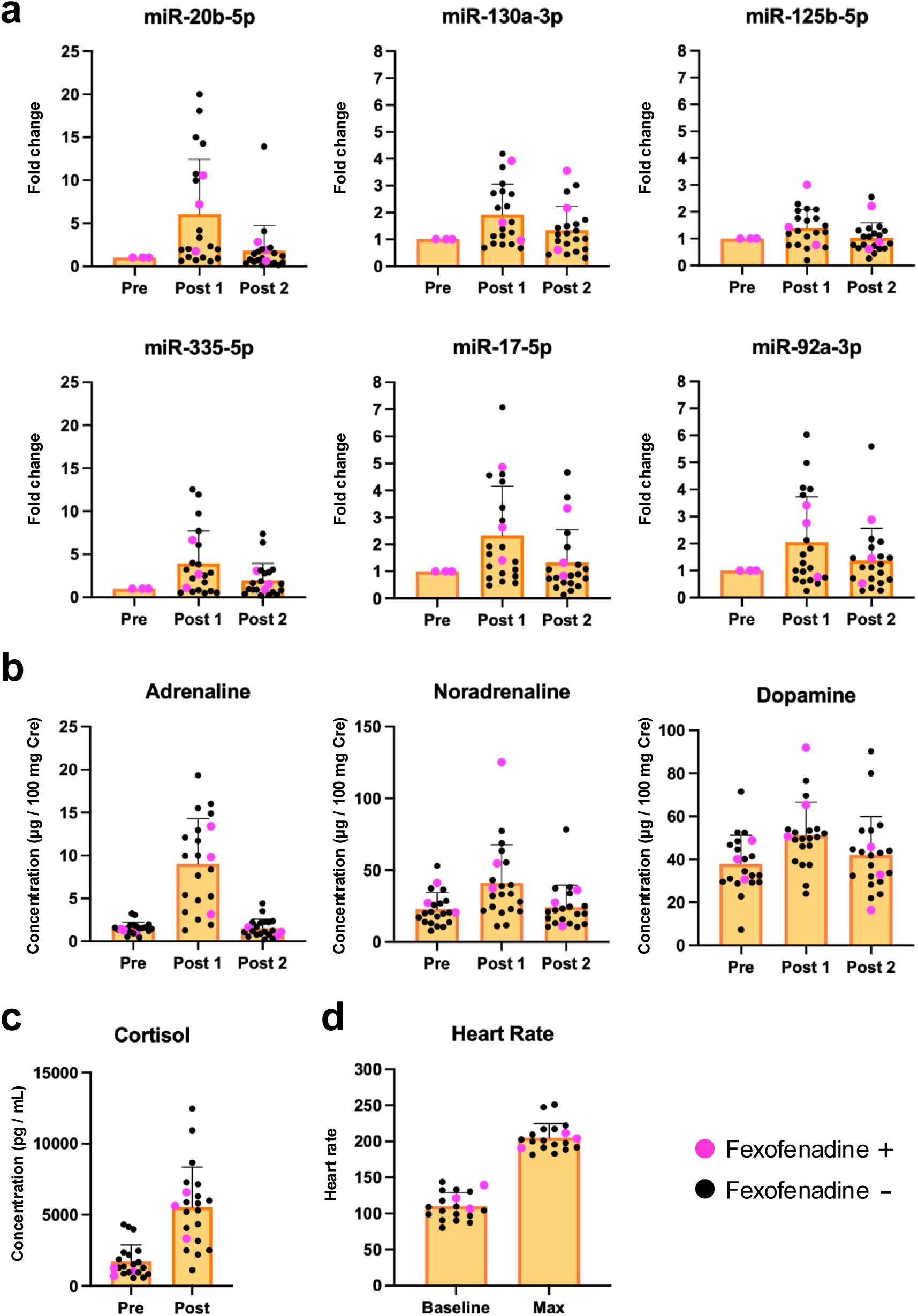
Fexofenadine had no effect on miRNA, cortisol, or catecholamine levels. **a**, microRNA. **b,** Catecholamine. **c,** Cortisol. **d,** Heart rate. Data are means±SD (n=21).

**Fig. S5.**
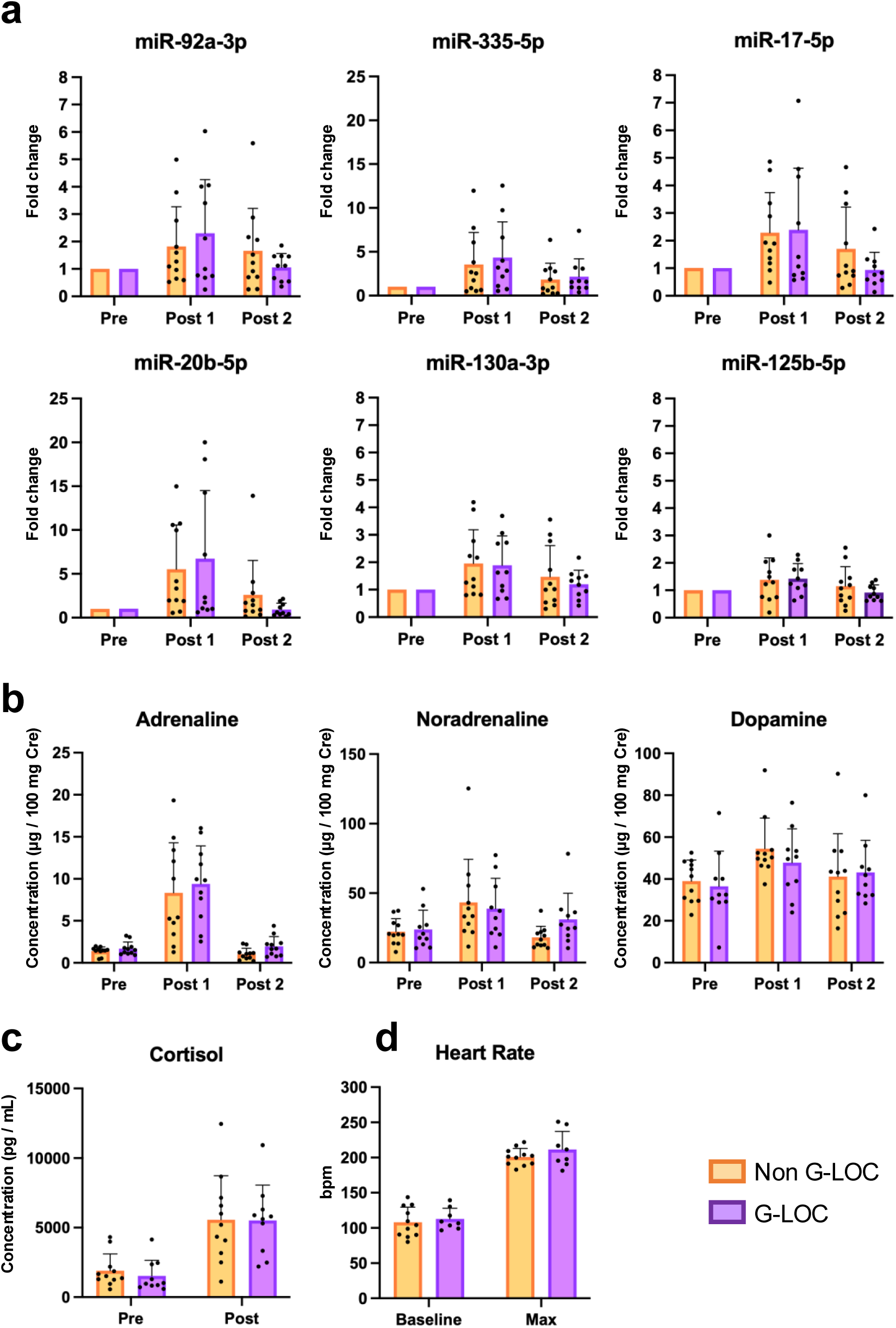
G-LOC did not alters uEV-encapsulated miRNAs and traditional stress hormones. **a**,**b**,**c,d** G-LOC did not affect the levels of six miRNAs, urinary catecholamines, salivary cortisol or heart rate, which were increased by +Gz stress (Figure 4a-e). Data are means±SD (n=21). Statistical analysis was performed using two-way ANOVA. G-LOC, gravity-induced loss of consciousness. uEV, urinary extracellular vesicles.

